# A deep learning-generated hASIC1a miniprotein inhibitor confers neuroprotection in stroke

**DOI:** 10.64898/2026.10.04.755611

**Authors:** Janina Sörmann, Konstantina Epitropaki, Maria Francesca Vicino, Christina Latt Fjorbak, Ciara Frances Pugh, Bernadette Lang, Emma Krindel Beyer, Raquel Comaposada-Baró, Fabian Hink, Søren Friis, Samuel Usher, Changsi Cai, Henriette Elisabeth Autzen, Joseph Matthew Rogers, Stephan Alexander Pless

## Abstract

Ion channels remain notoriously recalcitrant drug targets despite their extensive pathophysiological significance. Acid-sensing ion channels (ASICs) are proton-gated cation channels that cause neuronal death when excessively activated, such as during ischaemic stroke. This acidosis-driven injury pathway remains unchallenged by existing therapeutics. Here we generate two *de novo* designed hASIC1a-specific miniproteins by developing a platform that integrates computational miniprotein design, high-throughput automated patch clamp screening, biophysical and structural characterisation, and *in vivo* validation. We identify a subtype-selective hASIC1a inhibitor, Denasin1, which achieves near-complete inhibition with nanomolar potency. Denasin1 is disordered in solution but adopts the computationally predicted helix-turn-helix fold when bound to the extracellular acidic pocket of hASIC1a. Crucially, Denasin1 reduces infarct volume by > 50% in a murine model of ischaemic stroke. We thereby establish our *de novo* design platform as a route to develop potent and selective ion channel modulators.

## Introduction

Ion channels are key regulators of cellular excitability and intercellular communication. Acid-sensing ion channels (ASICs) are proton-gated cation channels that modulate neuronal excitability throughout the central and peripheral nervous systems, contributing to vital physiological processes such as synaptic transmission and sensory perception.^1^ In humans, four genes encode six ASIC subtypes (hASIC1a, 1b, 2a, 2b, 3 and 4), which assemble into homo- and heterotrimeric complexes. Of these, hASIC1a is the most widely expressed subtype, particularly in the central nervous system.^2^ Each monomer in the trimeric complex has a conserved architecture comprising a large pH-sensing extracellular domain (ECD) connected to two transmembrane helices that line a central ion-conducting pore (Fig. 1a). Proton binding induces conformational changes that open a cation-selective pore.^2,3^

**Figure 1:**
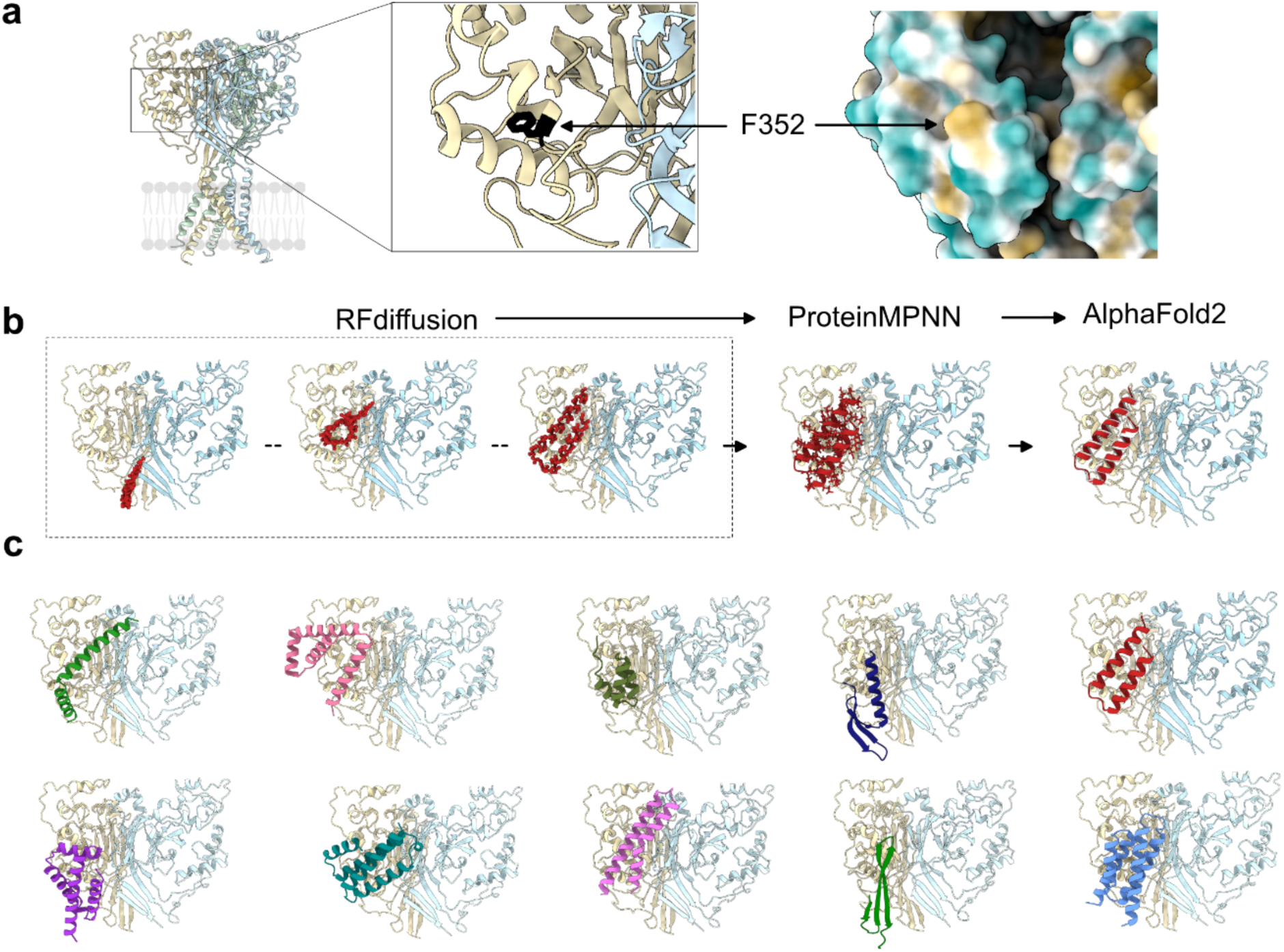
Computationally designed miniproteins converge on F352 as an interaction site. (a) Ribbon model of hASIC1a with colour-coded protomers (khaki, light blue, light green; left, PDB ID 7CFS). Insets highlight the position of F352 (black) in the ECD (left) and in hydrophobic surface representation (right). (b) Overview of computational pipeline. (c) Examples of *de novo* miniprotein designs with AF2-derived pAE scores < 14, highlighting the diversity of the predicted folds.

In pathological conditions such as ischaemia, inflammation or injury, persistent extracellular acidification causes sustained ASIC activation and eventually cell injury or death.^1^ Interestingly, genetic deletion or pharmacological inhibition of hASIC1a confers cardio- and neuroprotection.^4,5^ Although this highlights the importance of hASIC1a as a therapeutic target,^5–7^ effective ASIC1a inhibitors are lacking. Small molecule compounds such as NS383,^8^ amiloride, and amiloride derivatives ^9^ suffer from a lack of either selectivity or potency.^9–11^ Venom-derived peptide toxins, including Psalmotoxin1 (PcTx1) and Hi1a, show higher affinity and subtype-selectivity, decreasing infarct volumes and improving neurological and motor function in middle cerebral artery occlusion rodent stroke models.^12^ However, they display complex, highly pH-dependent pharmacology and their multiple reducible disulfide bonds limit large-scale production.^13–16^ Similarly, the antibody ASC06-IgG1 is a potent and selective inhibitor of hASIC1a,^17^ but the cost of production is relatively high and its size and complexity hinders tissue penetration. Together, these limitations underscore the need for a new class of hASIC1a modulators that combine nanomolar potency, subtype selectivity, and scalable production.

The challenge of discovering modulators is not unique to ASICs and reflects a broader challenge in ion channel pharmacology. Structural conservation within pore-forming and ligand-binding domains means that potent inhibitors of one subtype often exhibit off-target effects on related family members.^18^ Thus, despite the involvement of ion channels in numerous diseases and decades of pharmacological investigation, they remain recalcitrant to therapeutic intervention. Recent advances in *de novo* protein design have opened new opportunities to address these issues, enabling the generation of target-specific binding proteins with tuneable properties.^19,20^ In particular, deep learning methods that enable diffusion-based backbone generation and sequence optimisation have facilitated the computational design of miniproteins with tailored interaction geometries and defined binding surfaces at scale.^21,22^

In this study, we explored the potential of *de novo* protein design to generate ion channel inhibitors, using hASIC1a as a proof-of-principle. We created a streamlined end-to-end discovery platform integrating (i) computational design and scoring of candidate binders; (ii) high-throughput expression and purification; (iii) functional *in vitr*o screening using automated patch clamp (APC); (iv) biophysical characterisation; (v) structural validation; and (vi) *in vivo* testing. The platform identified multiple miniproteins with nanomolar potency against hASIC1a, the two most promising of which had high subtype-selectivity. Our approach generated a lead *<u>de n</u>ovo* <u>AS</u>IC1a <u>in</u>hibitor, Denasin1, which conferred robust neuroprotection in a mouse model of ischaemic stroke, outperforming an established venom-derived peptide inhibitor of hASIC1a.

## Results

### Computationally designed miniproteins converge on F352

To enable targeted modulation of hASIC1a, we first defined structurally and functionally relevant interaction sites within the ECD based on experimental closed-state structures (PDB ID: 7RNN, 7CFS, 7CFT). We focused on sites at or near the acidic pocket, a key determinant of proton sensing and channel gating, as well as a known drug- and toxin-binding site (Fig. 1a).^23–26^ Within this region, F352 was identified as a prominent accessible hydrophobic hotspot, located near the interface between each subunit (Fig. 1a). A range of hotspot combinations (F352 alone, F352 together with neighbouring residues, and combinations of residues elsewhere in the ECD) were used to guide binder placement during the diffusion-based design process.

We next implemented a three-step, deep-learning-guided design pipeline to generate miniprotein candidates targeting this site (Fig. 1b). Backbone scaffolds were generated using RFdiffusion^27^ with a fixed-structure ECD and hotspot constraints biasing interface formation. Amino acid sequences were assigned using ProteinMPNN,^28^ followed by structure-based filtering using AlphaFold2^29^ to evaluate folding confidence and predicted binding modes. Across all trajectories, we generated ∼12,000 backbone conformations and ∼70,000 sequence variants, which were filtered to 76 high-confidence candidates with predicted alignment error (pAE) scores below 14 for downstream experimental validation. Engagement of the F352 hotspot was observed in all high-scoring designs. Even runs in which hotspots were defined elsewhere in the ECD often resulted in high-scoring designs engaging F352, indicating pipeline-wide convergence on this interaction site.

The resulting miniprotein designs exhibited structural diversity. Most encompassed α-helical bundles, but two contained β-sheets and five had mixed α/β topology (Fig. 1c), demonstrating that the computational pipeline can generate diverse structural solutions against a single target site.

### APC facilitates identification of potent and selective inhibitors

To test whether any of the computationally predicted miniproteins could modulate hASIC1a function, we expressed and purified all 76 miniprotein designs in *E. coli* and applied them to HEK cells that express endogenous hASIC1a in APC experiments (Fig. 2). We recorded acidic pH-activated currents before and after incubation with a single-dose of each of the 76 miniproteins. Two initial miniprotein candidates, denoted as Denasin1 (44 amino acids) and Denasin2 (75 amino acids), showed > 50% inhibition. Denasin2 was predicted to form a four-helical bundle, while Denasin1 was predicted to comprise a helix-turn-helix motif (Fig. 2 a, d). Both were predicted to bind to a cluster of amino acid side chains around F352 near the hASIC1a acidic pocket. We confirmed hASIC1a engagement for both Denasin1 and Denasin2 using flow-induced dispersion analysis (FIDA) (Supplementary Fig. 1) before proceeding with them for further characterisation.

**Figure 2:**
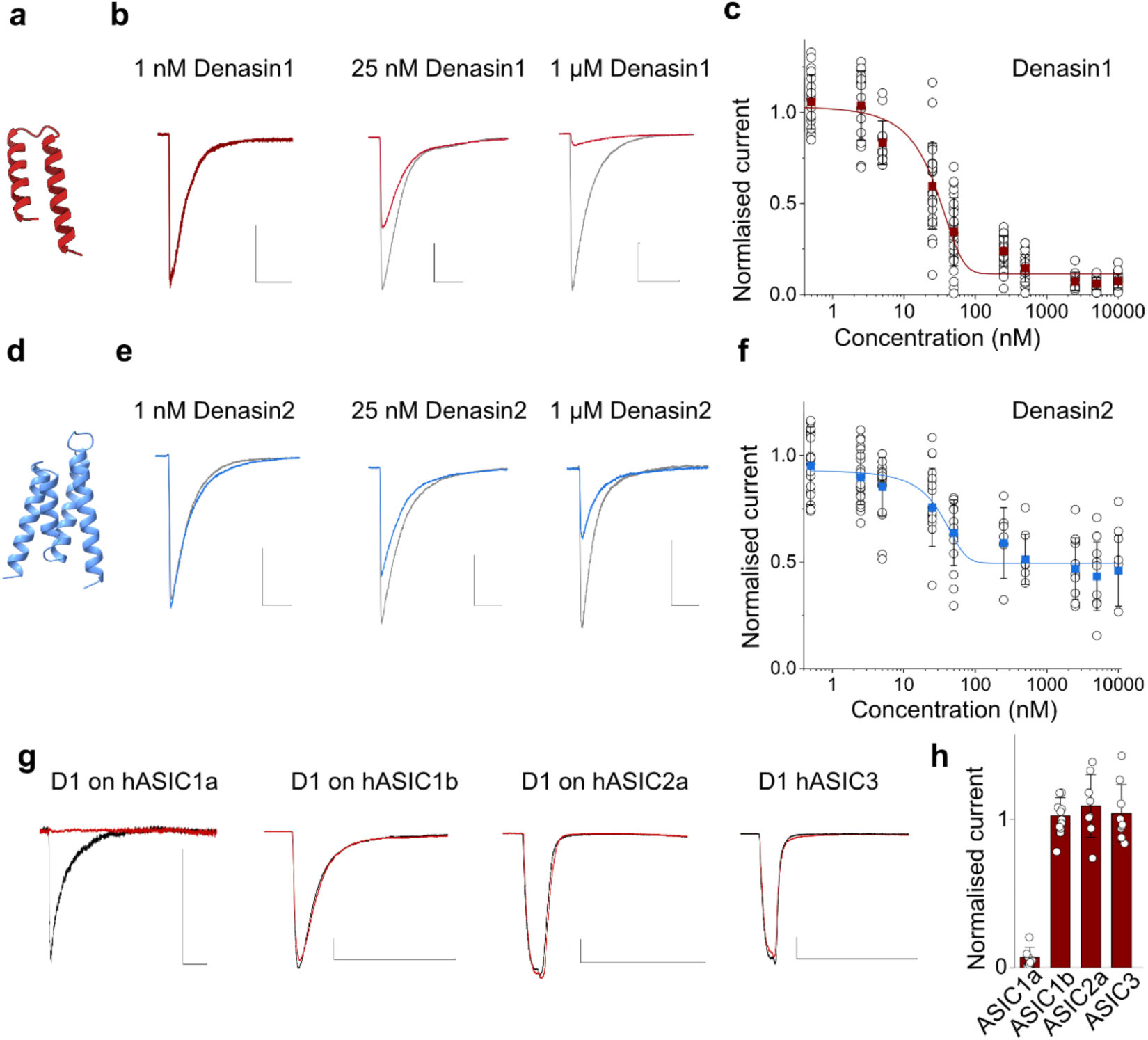
APC screen identifies potent and selective ASIC1a inhibitors. (a) Structural prediction of Denasin1. (b) Example pH-induced hASIC1a currents without (grey) or with (red) increasing concentrations of Denasin1. (c) Concentration–response curve for Denasin1. (d) Structural prediction of Denasin2. (e) Example pH-induced hASIC1a currents with Denasin2 (blue). (f) Concentration–response curve for Denasin2. (g) hASIC1a, hASIC1b, hASIC2a, and hASIC3 currents before (grey) and after (red) Denasin1 (D1) application. (h) Bar plot of currents after Denasin1 application normalised to currents before Denasin1 application. Scale bars represent 2 s (x-axis) and 1 nA (y-axis), but for Denasin1 on hASIC1a the scale bars represent 0.5 s (x-axis) and 1 nA (y-axis). Error bars represent S.D.

We expressed and purified Denasin1 and 2 at a larger scale and evaluated their potency and efficacy in APC recordings. Denasin2 inhibited hASIC1a currents with nanomolar potency (IC_50_ = 30 ± 5.1 nM) and an efficacy of ∼50% (Fig. 2d, e). Strikingly, Denasin1 caused near-complete inhibition of hASIC1a-mediated currents with an IC₅₀ of 23 ± 2.5 nM (Fig. 2b, c). The near-complete block by Denasin1 and rank order of Denasin1/2 inhibition was reproduced in two-electrode voltage clamp (TEVC) recordings from *Xenopus laevis* oocytes heterologously expressing hASIC1a (Supplementary Fig. 2), albeit with lower apparent potency, consistent with the notion that different expression systems and acidification protocols can affect ASIC IC_50_ values.^14,15,30^ We next assessed the subtype selectivity of Denasins using APC recordings from ASIC1a-knockout HEK cells expressing hASIC1a, hASIC1b, hASIC2a, or hASIC3 via baculovirus transduction.^31^ At a concentration of 5 µM, Denasin1 achieved almost complete inhibition of hASIC1a, but left other hASIC subtypes unaffected (Fig. 2g, h and Supplementary Fig. 3). Similarly, Denasin2 failed to inhibit hASIC3 currents, even at high (10 µM) concentrations (Supplementary Fig. 3). This indicated robust subtype selectivity despite the structural similarity among hASIC family members.

To test if diversification of Denasin1 and 2 would yield additional high-potency candidates we used either partial diffusion, whereby the backbones of Denasin1 and 2 were subjected to further noising and diffusion steps,^27^ or ProteinMPNN sequence design repeated at different sampling temperatures.^28^ The resulting designs were predicted to bind around the acidic pocket and retained contacts with the previously identified hotspot while sampling additional structural diversity. Although this increased hit rates and yielded a substantial number of additional *de novo* hASIC1a inhibitors, it did not produce designs with higher potency or efficacy than Denasin1 and 2 (Supplementary Fig. 4).

These results confirm that we have established a scalable, hotspot-driven design framework that connects generative modelling with experimental validation, enabling the identification of subtype-selective ion channel modulators with nanomolar potency.

### Denasin1 utilises a fold-upon-binding mechanism

Because Denasin1 is a relatively short miniprotein without a readily identifiable hydrophobic core, we used circular dichroism (CD) analysis to characterise the folding of Denasins in their unbound state. Denasin1 displayed a low (< 10%) helical content in solution, whereas Denasin2 retained > 95% helicity, even after heating to 95°C (Fig. 3a and Supplementary Fig. 5). This suggests that Denasin1 is largely unstructured in solution, similar to the endogenous hASIC1a peptide modulator big dynorphin.^31,32^ To assess whether Denasin1 undergoes folding-upon-binding ^33–35^ to adopt the predicted helix-turn-helix fold, we determined the experimental structure of the hASIC1a-Denasin1 complex.

**Figure 3:**
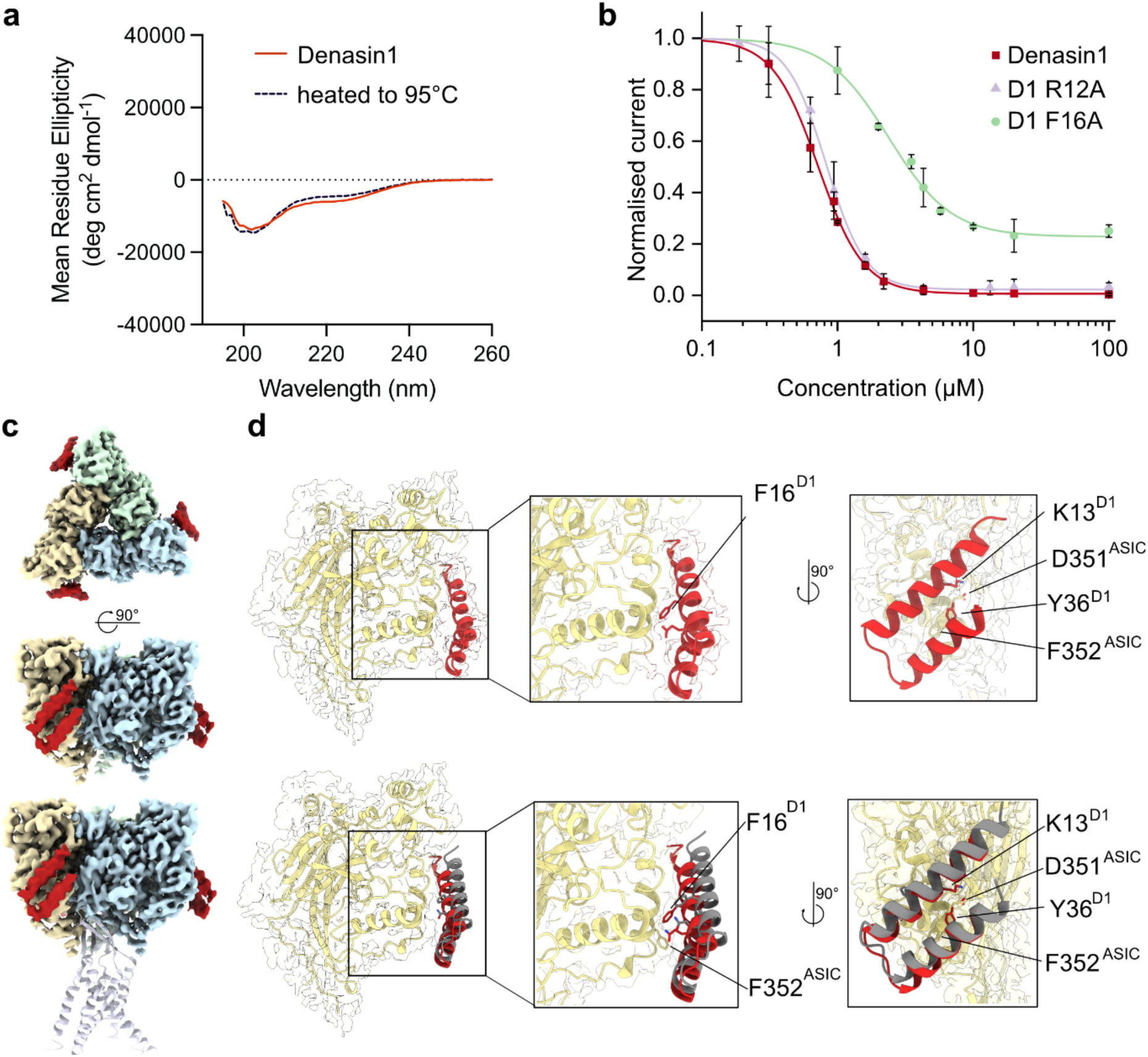
Denasin1 engages hASIC1a with a fold-upon-binding mechanism. (a) CD spectra for Denasin1 before (solid line) and after (dashed line) heating to 95C; (b) TEVC-derived concentration-response curves for Denasin1 (red) or R12A and F16A mutants (violet and light green, respectively) obtained with hASIC1a heterologously expressed in *Xenopus laevis* oocytes. Denasin1: 720 ± 12 nM (full block), R12A IC₅₀ = 820 ± 10 nM (full block), and F16A: IC₅₀ = 2.38 ± 0.12 μM and ∼80% block. (c) The unsharpened cryo-EM map of hASIC1a processed with C3 symmetry with protomers colored in khaki, mint green and light blue, respectively and Denasin1 in firebrick red as viewed from the side (top) and the extracellular side (middle). hASIC1a model (PDB ID 7CFS)^25^ represented in lavender cartoon is docked in to orient the cryo-EM map relative to the membrane plane (bottom). (d) Top: The sharpened cryo-EM map of hASIC processed with C1 symmetry for the single hASIC1a protomer (khaki) in complex with Denasin1 (firebrick red), with the corresponding atomic models of hASIC1a and Denasin1 shown as cartoon and with the side chains of P16 and Q20 or K13, Y36 (Denasin1), D351, and F352 (hASIC1a) shown in sticks. Bottom: Same as above, with the AlphaFold2 prediction of the hASIC1a-Denasin1 complex highlighted in gray.

We expressed and purified hASIC1a at large-scale, reconstituted it into membrane scaffold protein (MSP)-encircled nanodiscs (Supplementary Fig. 6), and determined its structure in complex with Denasin1 using single particle cryo-electron microscopy (cryo-EM, Supplementary Fig. 7,8,9, Supplementary Table 1). The 3D density map confirmed that Denasin1 adopts the predicted helix-turn-helix fold when bound to hASIC1a, exhibiting several well-defined contact points at the peptide-channel interface (Fig. 3c, d). The interface is centred around F352, which stabilises the interaction by engaging in an edge-to-face aromatic interaction with Y36 of Denasin1. Thus, although Denasin1 is largely unstructured in solution, it folds into its helix-turn-helix conformation upon engaging hASIC1a.

Our cryo-EM structure revealed that the hASIC1a-Denasin1 surface is predominantly hydrophobic, including F16 of Denasin1, which engages in a CH–π interaction with P348 of hASIC1a. By contrast, R12 of Denasin1 at the periphery of the interaction surface adopts two rotamer conformations, neither of which have well-defined interactions with hASIC1a. To confirm that side chains situated closer to the hydrophobic core of the experimentally determined binding interface make a greater functional contribution than those at the periphery, we mutated R12 and F16 in Denasin1 to alanine and measured the ability of these mutants to inhibit hASIC1a heterologously expressed in *Xenopus laevis* oocytes (Figure 3 b). Consistent with our structural observations and the notion that hydrophobic interactions drive the hASIC1a-Denasin1 interaction, R12A had no impact on inhibition whereas F16A significantly reduced both potency and efficacy, resulting in incomplete inhibition.

Together, these biophysical, structural and mutagenesis data confirm that Denasin1 binds to the computationally predicted binding site via a fold-upon-binding mechanism.

### Denasin1 confers robust neuroprotection *in vivo*

Due to its near complete inhibition of ASIC1a, as well as its smaller size, we prioritized Denasin1 for *in vivo* characterisation in a transient middle cerebral artery occlusion (MCAO) model of ischaemic stroke. Following pilot experiments to optimise occlusion duration and compound administration, we established a protocol of 30 min occlusion followed by 15 min reperfusion (Zea Longa model)^36^ before intrathecal administration of either Denasin1, vehicle (artificial cerebral spinal fluid, aCSF), or the established ASIC1a inhibitor PcTx1.^5^ Brains were extracted for analysis after 24 h (Fig. 4a). No increase in mortality or obvious adverse effects were observed during the experimental timeframe.

**Figure 4.**
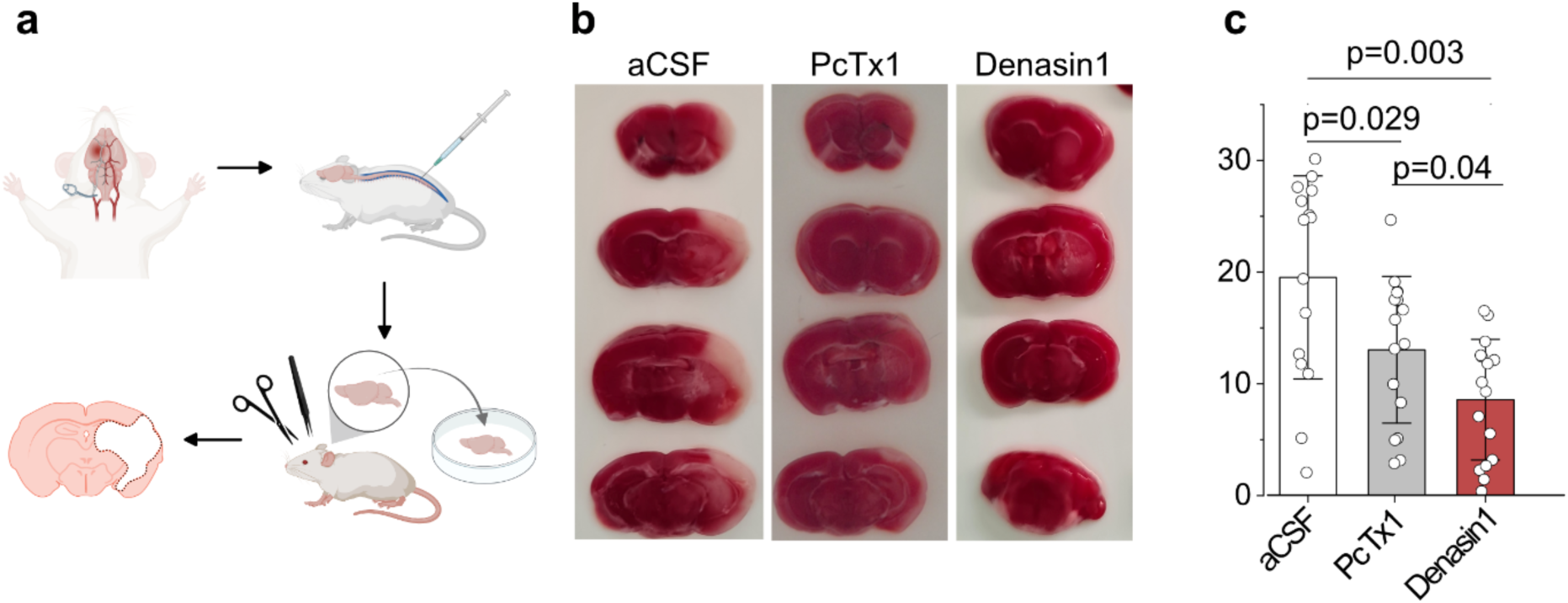
Denasin1 confers robust neuroprotection in a mouse model of ischaemic stroke. (a) Schematic overview of the MCAO stroke model (created in BioRender, Pless, S. (2026) https://BioRender.com/ wm050pe); (b) Representative TTC staining for brains treated with aCSF, PcTx1 or Denasin1. (c) Quantitative comparison of stroke volume after treatment with aCSF, PcTx1, or Denasin1. One-way ANOVA confirmed both treatments significantly reduced infarct volume compared to aCSF (PcTx1: p = 0.029; Denasin1: p = 0.003), with large (PcTx1, Cohen’s d = 0.82) and very large (Denasin1, Cohen’s d = 1.48) effect sizes (Supplementary table 2).

Strikingly, we observed a significantly lower infarct volume in Denasin1-treated animals compared to either aCSF or PcTx1 (Fig. 4b). Denasin1 reduced infarct size from 19.5 ± 9% (n = 16) in aCSF-treated animals to 8.6 ± 5.4% (n = 16) (Fig. 4c), exceeding that observed for PcTx1 (13.1 ± 6.6%, n = 16). Similarly, the infarct volumes after Denasin1 application were significantly smaller than after PcTx1 treatment (Supplementary Table 2).

Thus, although the established inhibitor PcTx1 confers significant neuroprotection *in vivo* relative to vehicle control, Denasin1 is more efficacious in the MCAO model under the tested conditions. Together, our results establish the translational potential of our platform for generating functional ASIC1a inhibitors.

## Discussion

We have developed a scalable and transferable platform to design *de novo* miniproteins targeting ion channels. Our approach integrates deep-learning-generated *de novo* miniprotein binder design with high-throughput expression, purification, functional screening, biophysical characterisation and *in vivo* validation. We demonstrate its utility by identifying a potent, subtype-selective ASIC1a inhibitor, Denasin1, which confers neuroprotection in the MCAO mouse model of stroke by reducing infarct volume significantly more than the known hASIC1a inhibitor PcTx1. Thus, our framework provides a springboard for the discovery of therapeutically relevant miniprotein modulators of previously intractable ion channel targets. The results show that precision biologics against ion channels can be accessed using computational design, laying the groundwork for next-generation therapeutics targeting the broader membrane proteome.

A prominent feature of our platform is its practical speed and efficiency. Critically, no intermediate quality control steps for expression yield or solubility are required before functional screening, and a binding screen is not necessary before electrophysiological assessment to specifically identify ion channel modulators. The identification of two functional inhibitors from the first round of just 76 candidates underscores the robustness of the computational filtering strategy and the efficiency of the integrated design–expression–test platform, demonstrating that deep learning-guided design can dramatically compress the hit identification phase in the discovery process. This contrasts with conventional, industry-standard high-throughput screening approaches, in which libraries containing millions of molecules must be screened to identify a lead, a time- and cost-intensive process.

Our miniprotein designs reflected the bias of RFdiffusion towards α-helical backbones,^37,38^ whose tightly packed geometries leave limited space to accommodate bulky hydrophobic residues. Accordingly, only two of the 76 first-round candidates contained tryptophan despite its potentially favourable contribution to binding interfaces. Remarkably, Denasin1 had one of the highest (least favourable) pAE scores among the tested designs, highlighting the limits of structural prediction scores for hit identification and the importance of experimental validation. Our first-round hit rate of ∼3% is consistent with the notion that identification of functional modulators is more challenging than identifying pure binders (note that our APC screen identifies only modulators). The approach is corroborated by the substantially higher hit rate observed in the second round of design following scaffold enrichment (32%, Supplementary Fig. 4), which compares favourably with the ∼19% success rate reported for binders of soluble targets using RFdiffusion.^27^

A central feature of our design work is the identification and utilisation of F352 within the ECD of ASIC1a. This residue, positioned within the acidic pocket and proximal to inter-subunit interfaces, emerged as a recurrent interaction hotspot across high-scoring miniproteins designs. The convergence of independent design trajectories on this site, together with the functional activity of resulting binders, supports a model in which engagement of F352 in the closed state of the channel contributes to stabilisation of non-conducting states. This region overlaps with known toxin-binding sites,^39–41^ suggesting that the pipeline recapitulates key structural determinants of ASIC1a modulation to design *de novo* scaffolds. Functional characterisation did not indicate any inhibition of hASIC1b, hASIC2a, or hASIC3 by our lead miniprotein Denasin1, revealing its remarkably high selectivity for hASIC1a under the conditions tested here. This is noteworthy, because some of the key amino acids around the acidic pocket, including F352, are conserved between hASIC1a and hASIC1b, highlighting the ability of our platform to achieve high subtype selectivity when targeting relatively conserved sites.

Interestingly, the CD spectrum for Denasin1 indicated that it is largely unstructured in solution, yet our cryo-EM co-structure confirmed its predicted helix-turn-helix fold when bound to hASIC1a in MSP lipid nanodiscs. Such fold-upon-binding behaviour is an abundant interaction mode between human proteins and does not preclude high affinity or selective interactions.^33–35^ Moreover, an unstructured miniprotein is not necessarily incompatible with *in vivo* activity. For example, the GLP-1 analogue Semaglutide is largely unstructured^42^ before it binds to its receptor in a helical conformation.^43^ We note that the size of Denasin1 (44 amino acids) lies in a range that is rarely explored for *de novo* binders as convention considers it too large for peptide discovery campaigns ^44–46^ and considered too small for miniproteins. Nonetheless, we have demonstrated that valuable *in vivo*-active molecules can be identified in this intermediate size range.

Our work provides a significant contribution to the nascent and rapidly evolving field of deep-learning-guided *de novo* miniprotein designs to target different membrane proteins. For example, Mahling *et al.* engineered a structure-guided peptide modulator of Na_V_1.5 inactivation, achieving sub-micromolar affinity and functional rescue in cardiac disease models, but the intracellular site of action presents a delivery barrier and the targeting of an endogenous protein-protein interaction site limits generalisation.^47^ A more recent study extended *de novo* miniprotein design to G protein-coupled receptors, describing selective agonists capable of distinguishing active from inactive receptor conformations, however *in vivo* efficacy has yet to be demonstrated.^21^ Together with our own work, these studies highlight how computational *de novo* design of miniproteins is rapidly emerging as a practical strategy to generate functional membrane protein modulators. Future advances in generative design and high-throughput screening will no doubt accelerate progress across the membrane proteome. Our platform is built with this trajectory in mind: a scalable, transferable framework designed not for a single target, but for broad application to electrogenic membrane proteins.

Despite these advances, our work has highlighted opportunities for further development. First, although our open-source computational pipeline produced a large number of high-confidence designs, only a subset exhibited functional activity, indicating that computational scores cannot be assumed to translate into modulation and therefore *in vivo* efficacy. Second, although both Denasin1 and 2 bind to and modulate hASIC1a, their different functional profiles caution that similar target regions can result in functionally distinct outcomes. In our case, this likely reflects the well-established complexity of ASIC gating, in which the allosteric coupling of ligand binding to channel opening are highly pH dependent and not necessarily proportional.^2,14,15,23,24^ Lastly, although we used intrathecal administration as a direct and controlled route to establish proof-of-principle *in vivo* efficacy for Denasin1, this approach was not intended to reflect a clinical delivery strategy. Future work will need to address the pharmacokinetic properties of Denasin1, including plasma and central nervous system half-life, stability in biofluids, toxicity, and blood-brain barrier penetration, to identify viable routes of administration for therapeutic development. With these developments, we anticipate that combining open-source design tools with high-throughput functional validation will be generalisable across ion channel families and have a major impact on the field. Furthermore, the modularity of the design and validation process renders this an ideal framework for accelerating the discovery of modulators for a wide variety of electrogenic membrane proteins for which experimental structures and APC-suitable cell lines are available.

## Methods

### Computational design pipeline for human ASIC1a binders

hASIC1a structures (PDB IDs: 7RNN, 7CFS, 7CFT) were used as templates. Structures were pre-processed by removing non-protein atoms and truncated to one or two subunits of the extracellular domain. A three-step design pipeline was implemented combining backbone generation, sequence optimisation and structure-based filtering. Binder backbones were generated using RFdiffusion with defined hotspot residues that guided the binder design towards the acidic pocket. Backbone lengths were constrained to 30–100 amino acids, and multiple independent diffusion trajectories with different hotspots were sampled for the different target preparations. Amino acid sequences were generated using ProteinMPNN, generating six sequence variants per backbone. Designs were filtered using AlphaFold2, evaluating both binder folding and complex formation. Candidates were ranked based on predicted local distance difference test (pLDDT) scores and interface predicted alignment error (pAE). Designs with pLDDT >85, interface pAE <14 Å, and structurally plausible binding modes without steric clashes were retained. In total, ∼12,000 backbones and 70,000 sequences were generated, of which 76 candidates were selected for experimental validation.

### Cloning and small-scale expression

*E.coli* codon-optimised genes coding for the 76 candidate miniproteins (ordered from Twist Bioscience) were cloned into a pET28-derived expression vector containing an N-terminal His_8_-tag and 3C protease cleavage site using Golden Gate assembly. Constructs were transformed into *E. coli* BL21(DE3) and grown overnight in LB media starter culture using a 96-well format. 100 µL of starter culture were added to 4 mL Overnight Express Instant TB Medium (Merck) in 24 deep well plate. They were incubated in a shaker at 37 °C for 5 h followed by 18 °C overnight. For purification, cells were centrifuged (4000 g for 10 min), treated with lysis buffer (50 mM Tris, 300 mM NaCl, 10 mM imidazole pH 8) and lysozyme with a final concentration of 0.25 mg/mL and stored at -80 °C for at least 1 h. After thawing, 10 µg/mL DNAse I (1 U/μL, Thermo Fisher Scientific) and 20 mM MgSO_4_ were added to the pellet and mixed. Debris was removed via centrifugation for 30 min at 13000 rpm at 4 °C. Miniproteins were purified via Ni²⁺-NTA affinity chromatography (Ni Sepharose™ 6Fast Flow, Merck on AcroPrep 24-well Filter Plate, 5 µm Supor Filter Plate, Cytiva). Miniproteins were eluted by on-column 3C protease cleavage directly into assay-compatible buffer. Expression and purification conditions had been optimised in preliminary experiments, and automated patch clamp served as the sole functional readout for all candidates.

### Expression and purification of miniproteins

Selected miniproteins (Denasin1, Denasin2, Denasin1.1, Denasin2.1 and Denasin2.2) were expressed and purified with the same approach in litre-scale *E. coli* BL21(DE3) cultures but eluted off the Ni²⁺-affinity chromatography (Ni Sepharose™ 6Fast Flow, Merck on Econo-Pac^®^ Chromatography Columns, Biorad) with 50 mM imidazole, followed by desalting chromatography on the ÄKTA system (HiTrap® Desalting Columns Cytiva) system or on a PD10 desalting column. Protein identity and purity were confirmed by UPLC and LC–MS.

Binder sequences were as follows:

Denasin1:

### SVEEQRKEKVERKIKFAKKQIASGAPPERQKKLEEYIEELKKKS

Denasin2: SMEKLRIEQRRLAIKIYSAIKLGTGTLEELKKLVDELASLDPELSLKLLEYLEEYAKKVAEEKKGSPLYPKYAEIAEYIAKQVEKLKKELEKL

Denasin1.1:

### AREEQRKEKAERKIKFAEKLIESGAPEEKAKKLKEYKKEVIEKS

Denasin2.1: EEEEKKKEAYKLFIKIYSAIKKGTGDLEELKKLVDELASLDKEYSLKLLKYLAEEAEKLAKEKKGSPIEKKLKELSKYIKEQVKKLEKELEEK

Denasin2.2: SSSEKRIELNRLTVEVVSGIKTGTATKEEIFAKIDEIASLDPEEGLKALEYINEYAKAEAKEKAGSPLYPILKAVAEYTQKKVDELKKKIEEK.

### UPLC

Purity was determined by ultra-high-performance liquid chromatography (UPLC) using a Waters Aqcuity system. Samples were analysed on a reverse-phase C18 column (Acquity UPLC BEH C18 1.7x50 mm, Waters) applying a linear gradient of 5-95% buffer B in buffer A over 7 minutes (A: H₂O/MeCN/TFA, 95:5:0.1%; B: H₂O/MeCN/TFA, 5:95:0.1%).

### LC-MS

Liquid chromatography mass spectrometry (LC-MS) was performed using an Agilent 6470 Quadrupole LC-MS system (Agilent Technologies) equipped with an AJS-ESI (Jet stream) ionization source. Samples were analysed on a reverse-phase C18 column (C-C18, 2.1x50 mm, 1.9 µM, Agilent Technologies) using a linear gradient of 5-95% buffer B in buffer A over 3 min. (A: H₂O/TFA, 99.9:0.1%; B: MeCN/TFA, 99.9:0.1%).

### Automated electrophysiology

Whole-cell recordings were performed using a SyncroPatch 384PE platform (Nanion). HEK293 cells were used for endogenous hASIC1a recordings, while ASIC1b, ASIC2a and ASIC3 were expressed using a BacMam system in hASIC1a knock-out HEK cells.^31^ Cells were voltage-clamped to −40 mV and stimulated by rapid extracellular acidification (pH 7.6 to pH 5.5). All experiments were performed on NPC S-type 4x chips, except for BacMam-transduced cells, for which NPC S-type 1x chips were used to accommodate the larger current amplitudes.

Baseline currents were recorded prior to binder incubation (300 s), followed by repeated activation. Inhibition was quantified as the ratio of post-treatment to initial currents. Each construct was tested across multiple wells and independent runs. Data were analysed using DataControl and OriginPro. Concentration response curves were corrected for run-down.

### Two-electrode voltage clamp (TEVC)

*Xenopus laevis* oocytes were injected with 5 ng cRNA encoding human hASIC1a and incubated at 18 °C for two days. For recordings, oocytes were voltage-clamped at −15 mV (OC-725C amplifier, Warner Instrument Corp.) and perfused with Ca²⁺-free ND96 solution (96 mM NaCl, 2 mM KCl, 1.8 mM BaCl2, 1 mM MgCl2, 5 mM HEPES pH 7.6) using an automated perfusion system (ValveBank, AutoMate Scientific). hASIC1a was activated by switching to Ca²⁺-free ND96 at pH 5.5. After two baseline activations, the miniprotein was applied for five minutes, followed by a 30 s wash in pH 7.6 buffer before two further activations, separated by a 1-minute interval. The average current amplitude of the two baseline activations was compared to the average of the two activations following binder application to quantify inhibition. Currents were analysed in Clampfit 11.3, filtered with an 8-pole Bessel low-pass filter. Current traces were exported to OriginPro 2020 for statistical analysis.

Mutant binders were generated by site-directed mutagenesis and tested under identical conditions.

### hASIC1a construct design and cloning

The codon optimized gene of human ASIC1a was purchased from Twist Bioscience and was cloned into a modified pEG BacMam^23^ vector using Gibson Assembly. The resulting construct contains an N-terminal 3X-FLAG tag followed by an eGFP-tag, separated from hASIC1a by a TEV protease cleavage site.

### hASIC1a expression

N-terminally GFP & 3x FLAG-labelled hASIC1a was expressed in HEK293S GnTi-cells which were seeded at an optical density of 1.0 x 10^6^ cells/mL in Freestyle medium supplemented with 2% fetal bovine serum (FBS) and incubated on an orbital shaker at 37°C, 135 rpm and 5% CO_2_. The cells were transduced using the baculovirus expression system.^48^ A final concentration of 10 mM sodium butyrate (NaBu) was added to the cells to enhance protein expression 20h post-transduction and the temperature was lowered to 30°C. Cells were harvested 72 h post transduction by centrifugation at 1500 rpm for 20 minutes and flash frozen in liquid nitrogen. For APC selectivity test hASIC1b, hASIC2a, and hASIC3 were expressed in ASIC1a-knockout HEK cells^31^ with the same protocol. Expression was confirmed via FSEC and APC.

### hASIC1a purification

Cells were thawed on ice and resuspended in 1 mL/g cells HBS buffer supplemented with cOmplete^TM^EDTA-free Protease Inhibitor Cocktail (HBS-PI) and resuspended cells were mixed with an equal amount of HBS-PI supplemented with 20mM DDM/ 4 mM CHS. Cells were solubilized for 1.5 h at 4°C on a tube roller and non-solubilized cell debris was removed by centrifugation at 26,000 × g and 4°C for 30 min. The supernatant was incubated for 1.5 hours with 0.5 mL of pre-equilibrated anti-FLAG resin (GenScript) at 4°C to allow binding of the FLAG-tagged protein. The resin was washed three times in a gravity column: The resin was washed sequentially with three 5-CV washes of 20 mM HEPES (pH 7.4), 150 mM NaCl, 0.5 mM DDM/ 0.1 mM CHS, the second wash supplemented with 5 mM MgCl₂ and 5 mM ATP to remove co-purifying ATPases. Finally, the protein was eluted with elution buffer containing 20 mM HEPES (pH 7.4), 150 mM NaCl, 0.5 mM DDM/0.1 mM CHS, and 300 μg/mL 3x -DYKDDDDK peptide (Genscript). For size-exclusion chromatography, the samples were filtered with a 0.22 μm filter and injected onto a Superose 6 Increase 10/300 GL column (Cytiva) pre-equilibrated with 20 mM HEPES (pH 7.4), 150 mM NaCl, 0.5 mM DDM/ 0.1 mM CHS, run at a flowrate of 0.25 mL/min. The combined fractions of the protein peak were taken for FIDA experiments.

### hASIC1a nanodisc reconstitution

For the reconstitution into MSP nanodiscs, hASIC1a was affinity purified as described above. 2.5μM of hASIC1a was mixed with soy polar lipid extract (Soy PLEx; Avanti Research) at a molar ratio of 1:800 and incubated for 1 h on ice. Afterwards, MSP1D1 was added in a molar ratio of 1:10 (hASIC1a:MSP), resulting in a final molar ratio of 1:10:800 hASIC1a:MSP:Soy PLEx. After addition of de-gassed and pre-wetted biobeads, the solution was incubated for 1 h, the beads were then exchanged, and the sample was incubated over night at 4°C. Biobeads were sedimented and the supernatant was incubated with pre-equilibrated GFP-nb resin for 2 h at 4 °C. The GFPnb resin was prepared in house by coupling GFPnb to the NHS-Activated Sepharose 4 Fast Flow resin (Cytiva) following their standard protocol. The hASIC1a containing nanodiscs were eluted from the beads by TEV protease cleavage for 2 h before further purification of the nanodiscs using a Superose 6 Increase 10/300 column equilibrated in 20 mM HEPES (pH 7.4) and 150 mM NaCl. The purified sample fractions were pooled and spin-concentrated to 0.96 mg/mL (UV-vis spectrophotometer 280 nm) before analysis by single particle cryo-EM. Reconstitution into MSP1D1 nanodiscs was required to obtain the complex; parallel attempts to resolve the hASIC1a–Denasin1 complex in GDN/CHS micelles or in copolymer-based native nanodiscs (MAASTY)^49^ did not yield a reconstruction.

### Cryo-EM sample preparation and data acquisition

For structural elucidation of the hASIC1a-Denasin1 complex, hASIC1a was purified in MSP1D1 nanodiscs containing SoyPLEx lipids, as described above. Denasin1 was added at a threefold molar excess relative to hASIC1a monomer, and 4 μL of the sample was added to a glow-discharged Quantifoil R1.2/1.3 300-mesh Au holey carbon grid (Quantifoil, Micro Tools GmbH). The cryo-EM grid was blotted for 3 s at 4 °C and 100% humidity and vitrified using a Vitrobot Mark IV (ThermoFisher Scientific). Cryo-EM data were acquired on a Titan Krios G2 cryo-transmission electron microscope equipped with a Thermo Scientific Selectris X energy filter and a Thermo Fisher Scientific Falcon IVi detector, operated at 300 kV. 5,120 movies were collected using EPU (Thermo Fisher Scientific) at a nominal magnification of 165,000×, corresponding to a calibrated pixel size of 0.728 Å, with a defocus range of 0.6 to 2.0 μm and a total electron exposure dose of 60 e^-^/Å^2^. Movies were saved as TIFF. Information on the data collection are included in Supplementary Table 1.

### Cryo-EM image processing

Cryo-EM data processing was performed in CryoSPARC v4.7.1.^50^ Collected movies were patch-based motion corrected, CTF estimated and manually curated to 4,831 micrographs in CryoSPARC Live v4.7.1. Three parallel particle picking pipelines were implemented: (1) Particles were picked using blob picker, from which two rounds of 2D classifications identified 2D class templates suitable for template-based picking, which underwent three rounds of 2D classifications. (2) Micrographs were denoised prior to blob picking and two rounds of 2D classifications. (3) Particles were picked using blob picker, followed by two rounds of 2D classifications. Consecutive *ab initio* reconstructions were performed from these particles, with two and three classes implemented. The resulting initial model and its particles were used for non-uniform (NU) refinement applying C3 symmetry. For all pipelines, junk detector was used prior to extraction, all particles were extracted with a 1.456 Å pixel size (2X binned), and duplicates were removed leaving 1,109,909 particles. These underwent iterative heterogeneous refinements using a single “good” class and three “junk” classes. Initially, the good class used was the 20 Å low pass filtered structure from pipeline (3). Following each iteration, the particles from the good class underwent *ab initio* reconstruction, from which the new initial model was used as input for the next iteration until the resolution stopped improving. This particle stack was then input for a final heterogeneous refinement step with two good classes, followed by *ab initio* reconstruction of the best class and NU refinement. The remaining particle stack was then unbinned (0.728 Å pixel size) and the particle orientations were rebalanced to remove over-represented views. Global CTF refinement and reference-based motion correction^51^ of the particle stack were carried out prior to NU refinement applying C1 or C3 symmetry, yielding a 2.64 Å and 2.33 Å reconstruction, respectively. However, the density corresponding to Denasin1 displayed low occupancy, requiring further classification to improve resolution. First, the particles were symmetry expanded and underwent local refinement masking a single protomer. The refined structure was classified in 3D using a focus mask of the Denasin1, classes with Denasin1 bound were chosen and used for final local refinement masking a single protomer. This resulted in a structure of the individual protomer of hASIC1a bound to Denasin1 (C1) at an estimated resolution of 2.7 Å from 129,961 particles. Next, to determine a co-structure of hASIC1a trimer bound to Denasin1, particles from the previous 2.33 Å reconstruction were classified in 3D with a focus mask that encapsulated the three Denasin1 binding sites of the ECD. This resulted in a single class with all Denasin1 binding sites occupied, from which NU refinement of the particle stack applying C3 symmetry resulted in a 2.73 Å reconstruction of the hASIC1a trimer-Denasin1 co-structure from 40,610 particles. Details on image processing and final reconstructions of the hASIC1a-Denasin1 protomer and trimer co-structures are included in Supplementary Fig. 8 and Supplementary Table 1.

### Model building and refinement

The resulting cryo-EM density maps and the amino acid sequence of Denasin1 were used as input for de novo model building with ModelAngelo.^52^ The generated Denasin1 model was combined with the extracellular domain (ECD) from a previously published hASIC1a structure (PDB ID: 7CFS)^53^ and docked into the cryo-EM density using ChimeraX.^26^ The composite model was initially refined using Namdinator.^54^ Following three rounds of refinement, the model of one of the three protomers was further optimized through manual refinement and iterative real-space refinement in Coot^55^ and Phenix,^56^ using a general restraints setup, resulting in the C1 model. For a C3 model, this protomer model was copied according to the C3 symmetry axis and also optimized through manual refinement and iterative real-space refinement in Coot and Phenix, using a general restraints setup. Both models, C1 and C3, were refined and finally validated via the validation statistics of MolProbity^57^ and Phenix.^56^ All figures were prepared using UCSF ChimeraX.

### Flow-induced dispersion analysis (FIDA)

All experiments were conducted on a FIDA Neo instrument (Fida Biosystems ApS, Copenhagen, Denmark) using light-emitting diode (LED) induced fluorescence detection with an extinction wavelength of 480 nm and an emission wavelength of >515 nm. Permanently coated capillaries were used throughout this study to reduce stickiness (75 μm inner diameter, 375 μm outer diameter, 1 m total length, and 84 cm length to detection window). The assay buffer was 20 mM HEPES buffer pH 7.4, 150 mM NaCl, 0.5 mM DDM/0.1 mM CHS. All measurements were done using the capillary mixing approach in triplicates. Denasin 1 and 2 were expressed and purified for the FIDA experiments as described above. Detection of binding required hASIC1a to be solubilised in DDM/CHS; parallel measurements with hASIC1a in GDN/CHS micelles or in copolymer-based native nanodiscs (MAASTY) showed no concentration-dependent change in hydrodynamic radius. All FIDA experiments were carried out using the standard coated capillary method, which consists of following automated steps necessary to acquire a single measurement: Each measurement consisted of three washing steps including a water injection (40 s, 3500 mbar), a 0.1% phosphoric acid injection (60s, 3500 mbar), and a second water injection (40 s, 3500 mbar) followed by analyte injection (20 s, 3500 mbar), indicator injection with or without analyte (10 s, 50 mbar), and a 240 s data-acquisition run at 400 mbar.

### Binding analysis of Denasin1 and 2 to hASIC1a

Binding screening against hASIC1a was performed in a capillary mixing (capmix) mode by loading different concentrations of purified Denasin1 or 2 in steps 4 and 6 of the sequence and 15 nM of GFP-labelled hASIC1a in step 5. The hydrodynamic radius R_h_ values were calculated by processing the Taylograms using the FIDA Software v3.1 (Fida Biosystems) with a Taylogram fraction of 65%. For each independent experiment, dose–response curves were generated by plotting R_h_ as a function of log[Denasin1 or 2]. Curves were fitted individually using a four-parameter nonlinear regression model (log(agonist) vs. response with variable slope) in GraphPad Prism (v10.2). Following fitting, data were normalized to the fitted top and bottom values of each individual curve, where the bottom value corresponded to the no-mini protein control. Normalized datasets were then averaged across independent experiments, and the mean ± S.D. was calculated. Final curves are presented as %Bmax versus log[Denasin1 or 2] (M). Independent experimental replicates were defined as measurements performed on separately handled aliquots derived from independent protein purifications (affinity chromatography followed by size exclusion chromatography).

### CD spectroscopy

Circular dichroism (CD) spectra and thermal melt curves were obtained with a JASCO J-1500 spectrophotometer. During measurements, the temperature was maintained at 25 °C using a circulating water bath. The miniprotein Denasin1 concentration was adjusted to a 0.25 mg/mL (32 *μ*M) in HEPES buffer (150 mM NaCl, 50 mM HEPES). Measurements were conducted in 2 mm quartz cuvettes, and buffer-only spectra were recorded and subtracted from the sample data. For thermal melt experiments the temperature was increased to 95 °C and subsequently decreased, with CD spectra continuously monitored throughout the process.

### Middle cerebral artery occlusion (MCAO) as *in vivo* stroke model

All animal experiments were approved by relevant authorities and conducted in accordance with ARRIVE guidelines. Adult male C57BL/6J mice (20–26 g) were used for this proof-of-concept study to allow direct comparison with a previous study employing the same single-sex design.^7,58,59^ The mice were randomly assigned to treatment groups (Denasin1, PcTx1, or vehicle) in a double-blind design, with surgeon and data analyst blinded to group allocation. A power analysis of the pilot study data determined the effect size and indicated that 16 mice per group would be required. Transient focal cerebral ischaemia was induced using the intraluminal filament (Longa model), with occlusion maintained for 30 min followed by reperfusion; cerebral blood flow reduction and recovery were confirmed by laser Doppler measurement. Denasin1, PcTx1 (both 250 µg/mL, 7 µL), or vehicle (7 µL) were administered intrathecally 15 min after reperfusion. Brains were collected 24 h after MCAO, sectioned into 2 mm slices, and stained with 2,3,5-triphenyltetrazolium chloride (TTC) for infarct quantification. Infarct volumes were quantified from digital images using ImageJ. For statistical analysis, data are presented as mean ± S.D. Statistical analyses including ANOVA and power analysis were performed using OriginPro2020 (Supplementary Table 2).

## Data Availability

The reconstructed maps are available from the Electron Microscopy Data Bank (EMDB) under accession codes EMD-59424 (hASIC1a processed in C3) and EMD-59425 (hASIC1a processed in C1). The atomic coordinates based on the reconstructions have been deposited in the Protein Data Bank (PDB) under accession codes PDB 33GB (hASIC1a in C3) and 33GC (hASIC1a in C1). The raw cryo-EM movies and particles stacks have been deposited on the Electron Microscopy Public Image Archive (EMPIAR) and accession codes will be made available before publication. Plasmids and other data that support the findings of this study are available from the corresponding authors. Source data for all main figures will be made available via Figshare.

## Code Availability

The code used for protein design was obtained from https://github.com/RosettaCommons/RFdiffusion.

## Acknowledgements

We thank Dr. Lesley Anson for helpful comments on the manuscript.

## Funding

The project received support from Independent Research Fund Denmark (3103-00127B to S.A.P, J.M.R, H.E.A.; 5284-00320B to S.A.P and J.M.R.; 3105-00232B to J.M.R.), the Novo Nordisk Foundation (NNF20OC0060692 to H.E.A.; NNF25OC0100883 to J.M.R.), the Carlsberg Foundation (CF23-1153 to S.A.P and J.M.R.; CF20-0533 and CF23-0832 to H.E.A.), the Lundbeck Foundation (R480-2024-894 to J.S., R345-2020-1440, R518-2025-934 to C.C.) and the Hørslev Foundation (to J.S.).

## Author contributions

J.S., S.A.P., H.E.A., J.M.R. and C.C. conceived and designed elements of the study. S.U. installed RFdiffusion on the HPC cluster; J.S. adapted the protocol for hASIC1a binders and generated all miniprotein designs. J.M.R. provided a protocol for miniprotein expression, which J.S., S.U. and F.H. optimised. J.S. and M.F.V. expressed and purified the miniproteins. J.S. and S.F. designed the APC experiments, and J.S. performed and analysed the APC recordings. K.E. performed and analysed the FIDA experiments with guidance by H.E.A. J.M.R. performed and analysed the CD spectroscopy experiments. M.F.V. and C.F.P. performed and analysed the cryo-EM experiments with guidance by H.E.A. J.S. designed the mutational studies and TEVC experiments, which B.L. performed and analysed. E.K.B. performed and analysed the mass spectrometry experiments. K.E. and J.S. produced the BacMam viruses and transduced HEK cells. K.E. and M.F.V. performed large-scale expression and purification of hASIC1a with guidance by H.E.A. J.S. designed and analysed the MCAO study, which C.L.F. and R.C.B. carried out. J.S. and S.A.P. drafted the manuscript, and all authors critically revised it and approved the final version.

## Ethics declaration

All animal procedures were approved by the Danish National Committee on Health Research [Dyreforsøgstilsynet, 2021-15-0201-01101] and conducted in accordance with institutional guidelines and the European Council’s Convention for the Protection of Vertebrate Animals, consistent with the National Institutes of Health Guide for the Care and Use of Laboratory Animals. This study was conducted and reported in accordance with the ARRIVE guidelines.

## Competing interests

The authors declare no competing interests.

## Notes

### Competing Interest Statement

The authors have declared no competing interest.

