## Supplementary Information for "A deep learning-generated hASIC1a miniprotein inhibitor confers neuroprotection in stroke"

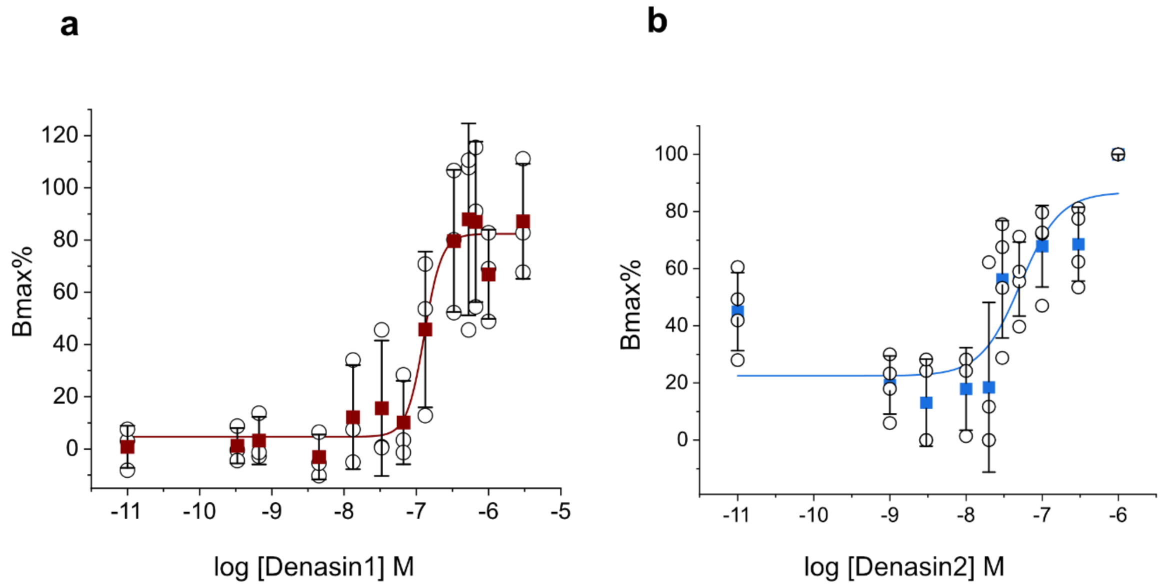


**Supplementary Figure 1:** Flow-induced dispersion analysis confirms binding. Denasin1 (a) and 2 (b) exhibited concentration-dependent binding to hASIC1a with affinities of 128 ± 52 nM and 55 ±16 nM.

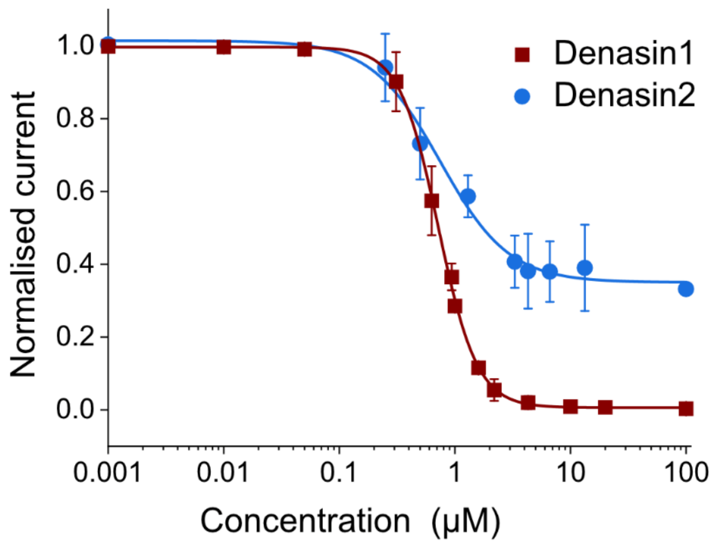


**Supplementary Figure 2:** Concentration-response-curves for Denasin1 and 2 obtained with TEVC. Denasin1 shows full inhibition and an $IC_{50}$ value of 720 ± 10 µM. Denasin2 shows 65% inhibition and an $IC_{50}$ value of 747 ± 115 µM.

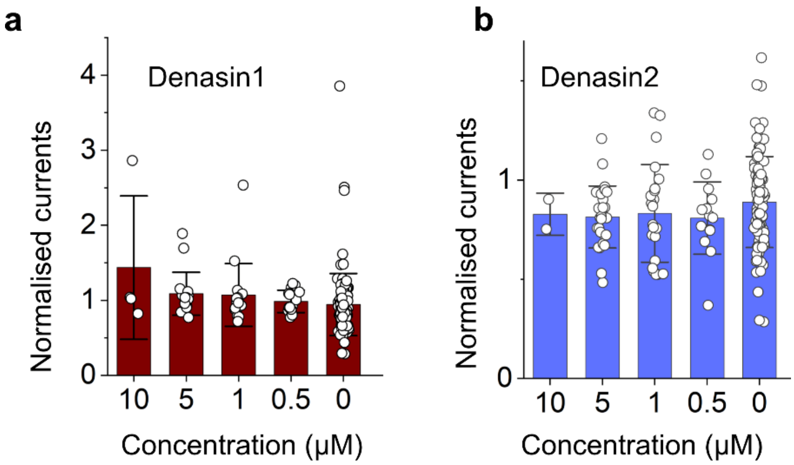


**Supplementary Figure 3:** Denasin1 and 2 do not inhibit hASIC3 in micromolar range. Concentration-response curve of Denasin1 (a) and 2 (b) on hASIC3.

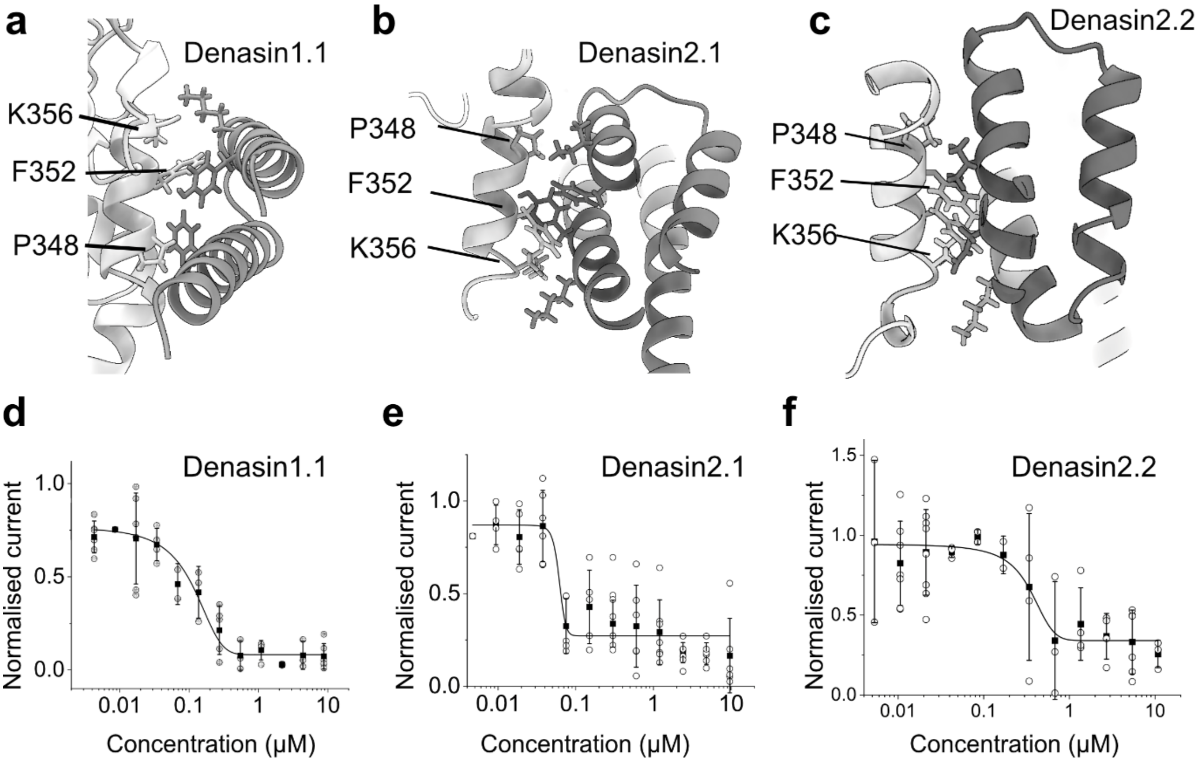


**Supplementary Figure 4:** Concentration–response curves and predicted binding poses of representative partial diffusion-derived binders. (a) Denasin1.1 shows an $IC_{50}$ value of 81.73 ± 13.83 nM. (b) Denasin2.1 shows an $IC_{50}$ value of 61.3 ± 19.24 nM. (c) Denasin2.2 shows an $IC_{50}$ value of 341 ± 52.99 nM. Denasin1.1, Denasin2.1 and Denasin2.2 are predicted to interact with residues P348, F352, and K356 of hASIC1a (PDB ID 7CFS).

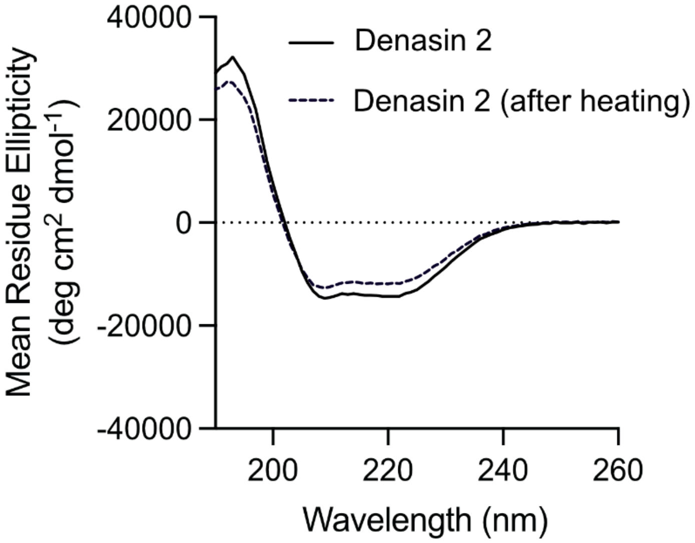


**Supplementary Figure 5.** CD spectrum of Denasin2. Circular dichroism (CD) spectra for Denasin2 before (solid line) and after heating to 95C (dashed line).

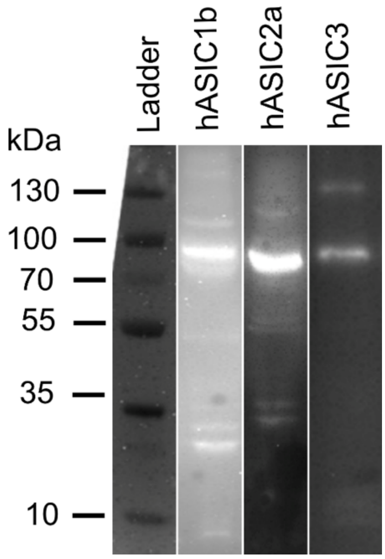


**Supplementary Figure 6:** Westen blots of Baculovirus-transduced hASIC1a KO HEK293 cells expressing hASIC1b, hASIC2a, or hASIC3. Proteins were detected using an anti-FLAG antibody.

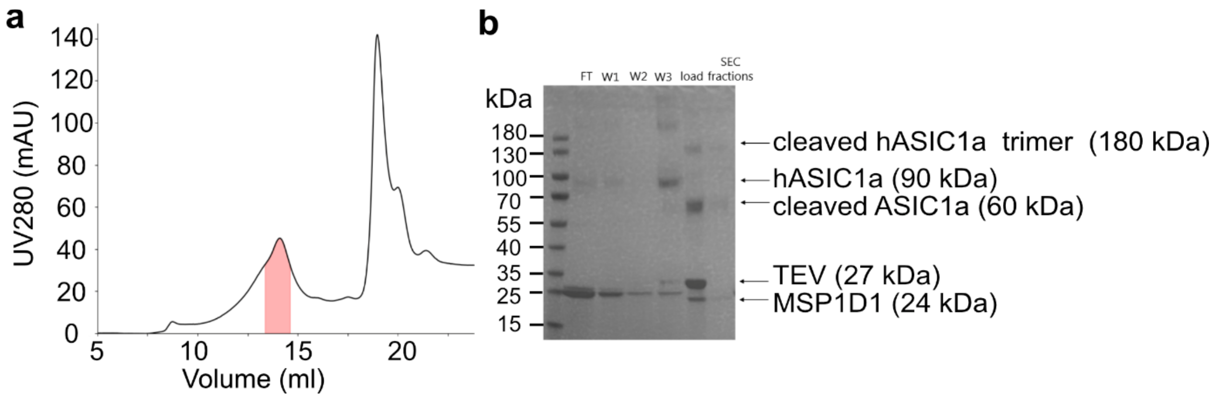


**Supplement Figure 7. hASIC purification for cryo-EM** (a) The size-exclusion chromatography (SEC) profile of hASIC1a in MSP1D1, where the highlighted fractions were used for cryo-EM sample preparation. (b) The SDS-Page analysis of the purification workflow. The gel shows the flow-through (FT) from GFPnb affinity chromatography, three wash fractions, and the sample loaded onto SEC, and the pooled SEC fractions used from cryo-EM sample preparation.

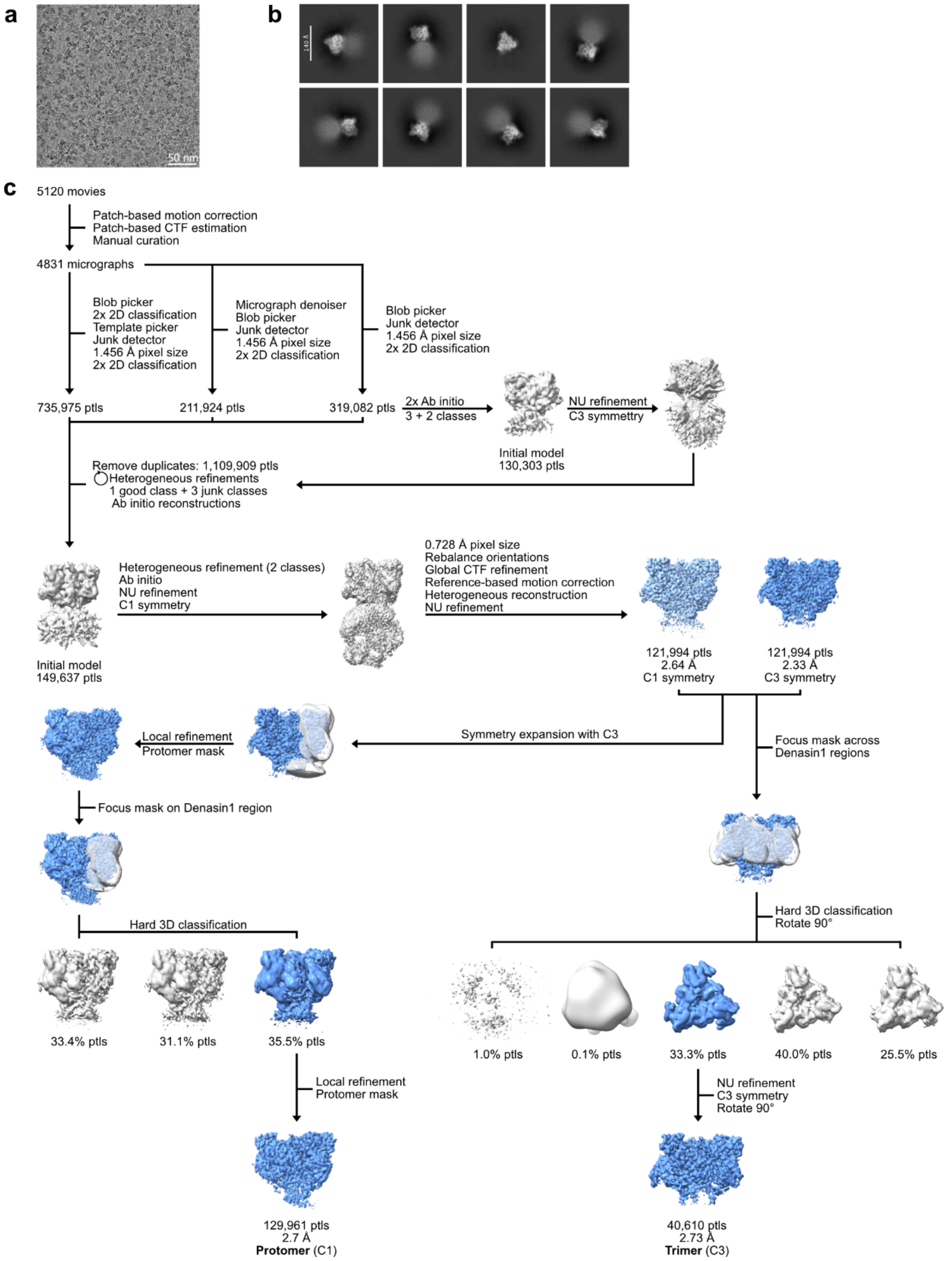


**Supplementary Figure 8:** Cryo-EM data processing of hASIC1a-Denasin1 co-structure in MSP lipid nanodiscs. (a) Representative motion-corrected micrograph from the dataset. (b) Representative 2D class averages. (c) Cryo-EM data processing pipeline to determine the hASIC1a-Denasin1 co-structure in MSP lipid nanodiscs. All data processing steps were carried out in CryoSPARC.[49] The data were motion-corrected, CTF estimated and manually curated before particle picking using multiple pipelines. Classification of the picked particles was carried out through iterative heterogenous refinements and *ab initio* reconstructions, with the final particle stack undergoing orientation rebalance, global CTF refinement and reference-based motion correction before non-uniform (NU) refinement. The refined particles were further classified to improve the resolution of the Denasin1 binding site using two methods: (Left) The particle stack was symmetry expanded applying C3 symmetry and particles were aligned to a single protomer using a mask. The particles were classified in 3D with a focus mask on the Denasin1 binding site resulting in a single class displaying Denasin1 occupancy, which was further refined resulting in a 2.7 Å resolution structure of hASIC1a protomer bound to Denasin1. (Right) A focus masked encapsulating all three Denasin1 binding sites was used for 3D classification of the particle stack, identifying a single class with all Denasin1 binding sites occupied. These particles were further refined by NU refinement applying C3 symmetry resulting in a 2.73 Å resolution structure of the hASIC1a trimer bound to three Denasin1 miniproteins.

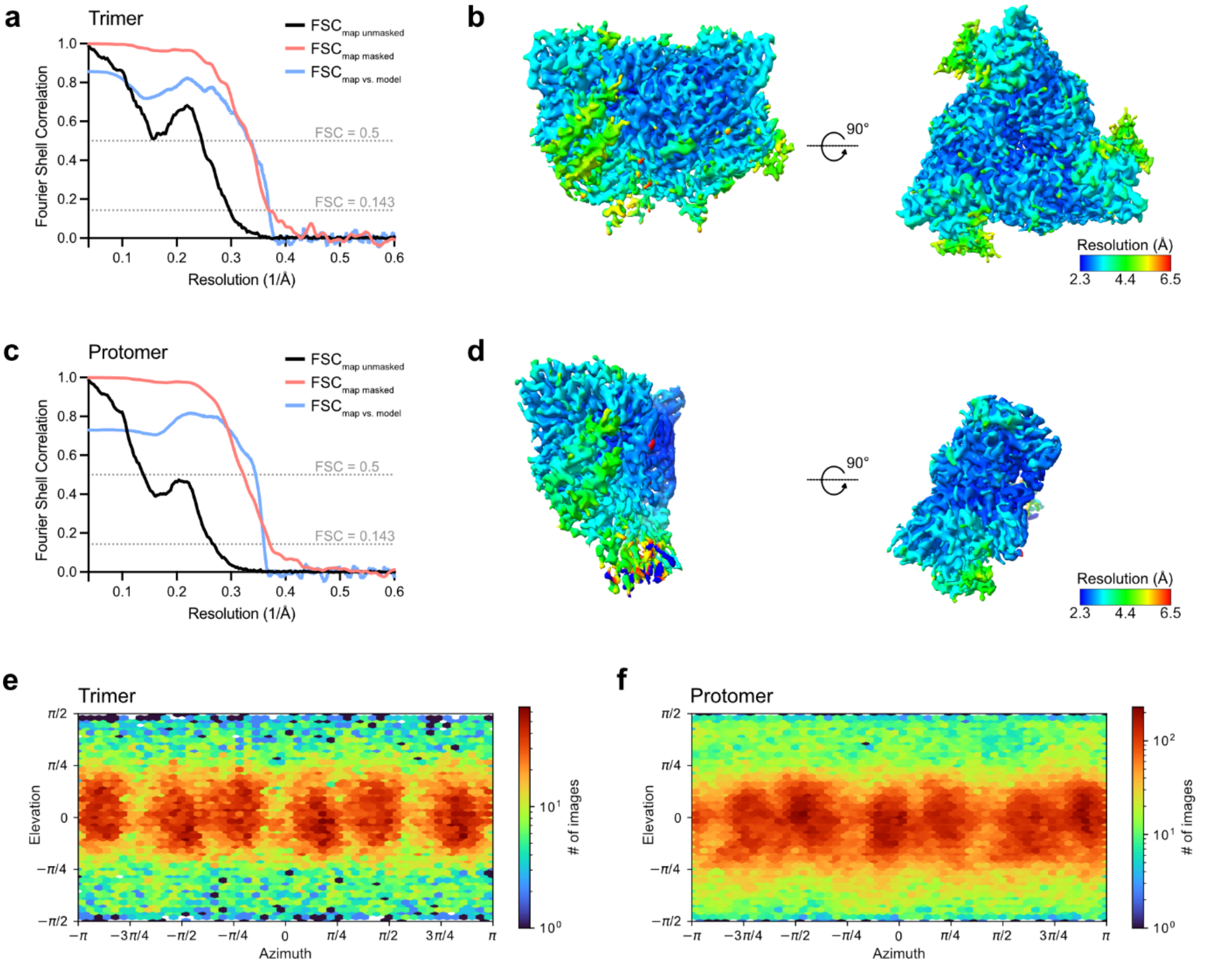


**Supplementary Figure 9:** Cryo-EM map and model validation curves, local resolution estimates and orientation distributions. (a) Fourier shell correlation (FSC) half map and model-map curves for hASIC1a trimer bound to Denasin1 (reconstructed with C3 symmetry) calculated by CryoSPARC[49] and Phenix Mtriage,[50] respectively. Gold standard FSC curves from the two half maps, $FSC_{map\ unmasked}$ (black) and $FSC_{map\ masked}$ (red). The $FSC_{map\ masked}$ curve utilises the automatically generated tight mask from CryoSPARC. The resolutions at FSC = 0.143 (dotted line) are 3.4 Å and 2.7 Å for $FSC_{map\ unmasked}$ and $FSC_{map\ masked}$, respectively. The $FSC_{map\ vs.\ model}$ curve (blue) is used for model validation, indicating a resolution of 3.0 Å at the FSC = 0.5 (dotted line) criterion. (b) Local resolution representation of the hASIC1a trimer-Denasin1 co-structure. (c) FSC half map and model-map curves for hASIC1a protomer bound to Denasin1. Descriptions are as in (a). The resolutions of $FSC_{map\ unmasked}$ and $FSC_{map\ masked}$ at FSC = 0.143 are 3.7 Å and 2.7 Å, respectively. The resolution of $FSC_{map\ vs.\ model}$ at FSC = 0.5 is 2.9 Å. (d). Local resolution representation of the hASIC1a protomer-Denasin1 co-structure. (e) and (f) Particle orientation distribution plots of the final refined hASIC1a-Denasin1 co-structures for the trimer and protomer, respectively.

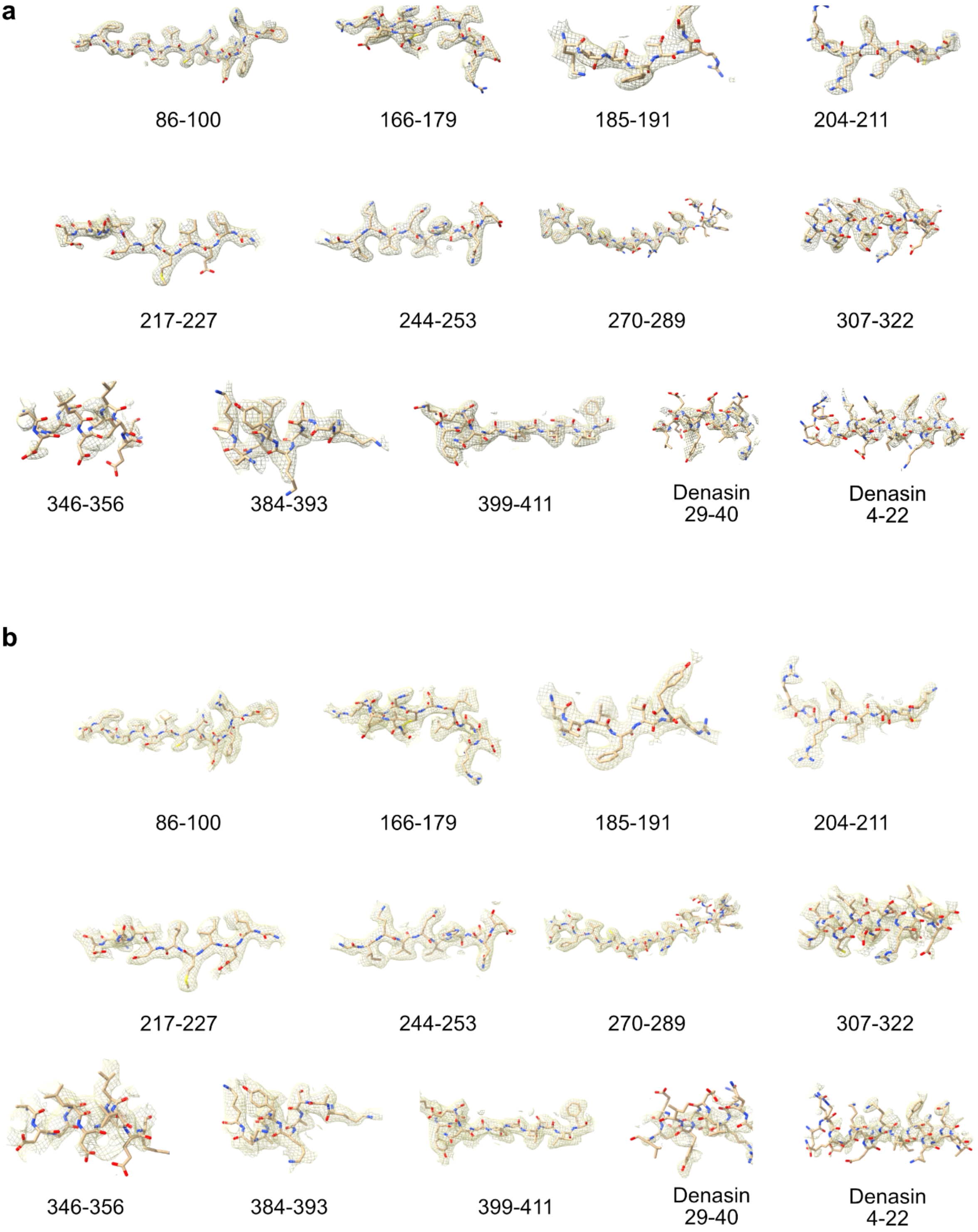


**Supplementary Figure 10:** Representative regions from the best reconstruction of the hASIC1a-Denasin1 complex in MSP nanodiscs and reconstructed in C1 and C3. Shown are representative labelled segments of hASIC1a ECD and Denasin1 with the fitted atomic model in C1 (a) and C3 (b).

**Table S1.** Data acquisition and processing, modelling statistics and validation scores.

| **Model refinement** | | |
|---|---|---|
| Protein residues | 71-423 (hASIC1a), 3-41 (Denasin) | 71-423 (hASIC1a), 3-41 (Denasin) |
| Ligands (pr chain) | 1 Denasin1 | 1 Denasin1 |
| **R.M.S. deviations** | | |
| Length (Å) | 0.004 | 0.003 |
| Angles (°) | 0.663 | 0.578 |
| **Ramachandran** | | |
| Favoured (%) | 94.33 | 93.99 |
| Allowed (%) | 5.67 | 6.01 |
| Outliers (%) | 0 | 0 |
| **Model validation** | | |
| MolProbity score | 1.83 | 1.88 |
| Clash score | 8.15 | 8.95 |
| Rotamer outliers (%) | 0.9 | 0 |
| Rama-Z score | -0.03 (0.45) | -0.74 (0.24) |

**Supplementary Table S2.** Data analysis of MCAO study. Statistical analysis using one-way ANOVA confirmed that both treatments significantly reduced infarct volume compared to vehicle (PcTx1: P = 0.029; Denasin1: P = 0.003), with large (PcTx1, Cohen's d = 0.82) and very large (Denasin1, Cohen's d = 1.48) effect sizes, indicating substantial biological efficacy. Furthermore, the infarct volumes after Denasin1 application were significantly smaller than those after PcTx1 application (P=0.044). Data was not corrected for edema.

| Compound | Infarct Volume [%] | Significance vs Buffer | Effect size Cohen's d | Significance vs PcTx1 |
|---|---|---|---|---|
| aCSF Buffer | 19.527 | | | |
| PcTx1 | 13.049 | P = 0.029 | (large) 0.82 | |
| Denasin1 | 8.59 | P = 0.003 | (large) 1.48 | P=0.044 |

To include mice into the study and into the analysis, the following criteria needed to be fulfilled:

- Mouse weight: 20-26 g
- Tail flick during intrathecal injection
- No topological bleeding
- No major core bleeding
- 70% reduction in laser doppler reading
- 70% reduction in the reading of laser doppler flow meter

In total, 63 mice were included, i. e., 16 mice/ compound group and 15 mice for aCSF group.
